# Sleep initiation difficulties involve weaker neural and physiological sleep transitions, particularly in children with neurodevelopmental conditions

**DOI:** 10.64898/2026.03.14.711131

**Authors:** Micha Hacohen, Miriam Guendelman, Ilan Dinstein

**Affiliations:** Psychology Department, Ben Gurion University of the Negev, Israel; Azrieli National Centre for Autism and Neurodevelopmental Research, Ben Gurion University of the Negev, Israel

## Abstract

The transition from wake to stable sleep is characterized by multiple neural, physiological, and behavioral changes. How these changes may differ in individuals with difficulties falling asleep such as children with neurodevelopmental conditions is poorly understood. Here, we studied sleep initiation in >2000 nights recorded from 186 children who participated in the Simons Sleep Project (SSP). Data included simultaneous, synchronized recordings of actigraphy, electroencephalography (EEG), photoplethysmography (PPG), and skin temperature. We extracted multiple neural, physiological, and behavioral measures that are known to increase/decrease during the sleep initiation period including EEG delta (1-4Hz) power, movement counts, heart rate (HR), and skin temperature. Transitions from 20 minutes before sleep onset to 40 minutes after sleep onset were modeled with a sigmoid function enabling the quantification of transition timing, speed, and magnitude per measure. Individuals with longer sleep onset latencies (SOL) exhibited smaller increases in EEG delta power and skin temperature as well as smaller decreases in HR and activity counts. These findings indicate that difficulties falling asleep are associated with multiple forms of cortical, physiological, and behavioral hyperarousal that can be measured at home with wearable devices. Importantly, transition magnitudes were key to explaining differences in SOL across participants (26% explained variance) in contrast to transition speed or timing within the sleep initiation period (<13% explained variance). Longer SOL and weaker transitions were particularly prominent in children diagnosed with autism and/or attention deficit hyperactivity disorder (ADHD).

## Introduction

Sleep onset is formally identified by visual inspection of electroencephalogram (EEG) recordings, according to American Academy of Sleep Medicine (AASM) guidelines, as the first 30-second epoch with sleep EEG. This epoch usually contains N1 stage sleep, which is defined as a noticeable reduction in alpha (8–13 Hz) power, increase in theta (4–7 Hz) power, emergence of low-amplitude mixed-frequency activity, slow eye movements, and a decrease in chin muscle tone (Troester et al., 2023). EEG characteristics indicative of sleep onset can differ across individuals (e.g., across ages) (Grigg-Damberger et al., 2007; Miraglia et al., 2024), making it difficult to reliably identify the precise epoch of sleep onset. This difficulty is evident, for example, in weak inter-rater reliability across expert sleep technicians when scoring N1 sleep epochs (Lee et al., 2022; Rosenberg & Hout, 2013). The transition from wake to sleep may, therefore, be more reliably characterized as a progressive process (Ogilvie, 2001), that includes a reduction in alpha power and increase in delta power (0.5–4 Hz), which take at least several minutes and reflect reduced flow of sensory input from the thalamus to the cortex and strengthening of widespread slow oscillations (slow wave activity – SWA) that characterize stable sleep (Borb & Achermann, 1999; Dijk & Von Schantz, 2005; Magnin et al., 2010).

The transition into stable sleep is also characterized by multiple physiological and behavioral changes. First, increased parasympathetic drive causes a decrease in heart rate and increase in heart rate variability (Boudreau et al., 2013; Chouchou & Desseilles, 2014; De Zambotti et al., 2011; Ma et al., 2024; Shinar et al., 2006). Second, skin temperature increases, particularly in the hands and feet, due to peripheral vasodilation that facilitates heat loss and enables the reduction of core body temperature (Kräuchi & Wirz-Justice, 2001a; Wyatt et al., 1999; Zulley et al., 1981). This process is generated by evening melatonin secretion, a hallmark of circadian sleep regulation (Kräuchi & Wirz-Justice, 2001a). Third, body movements decrease, a hallmark of sleep that is commonly monitored with accelerometers in actigraphy studies (Sadeh et al., 2000)

Difficulties transitioning from wake to sleep, measured by sleep onset latency (SOL) of > 30 minutes, is a diagnostic feature of Insomnia, a common sleep disorder affecting 10–15% of the population (Morin & Benca, 2012; Roth, 2007). Insomnia is particularly common in individuals with autism and/or attention deficit hyperactivity disorder (ADHD), which are two highly prevalent and closely related neurodevelopmental conditions (Richdale & Schreck, 2009). Actigraphy studies have reported that 60-80% of children and adolescents within these populations exhibit SOL of more than 30 minutes (Brevik et al., 2017; Øyane & Bjørvatn, 2005; Sung et al., 2008; Wajszilber et al., 2018). Prolonged SOL has been shown to exacerbate daytime behavioral and cognitive challenges (Bioulac et al., 2021; Knight & Dimitriou, 2019; McCabe et al., 2025), thereby motivating further research into the underlying mechanisms of sleep onset difficulties specifically in these clinical populations (Lord, 2019).

A leading theory of insomnia suggests that prolonged SOL may be caused by multiple interacting physiological, cognitive, and emotional regulation mechanisms that generate hyperarousal and interfere with the ability to initiate sleep (Bonnet & Arand, 2010). Potential mechanisms include inherent cortical hyperarousal evident in elevated high frequency power (Fernandez-Mendoza et al., 2017; Saper et al., 2010) that may interfere with the transition into widespread low frequency cortical synchronization necessary for sleep (Shi et al., 2022). Sensory over-responsiveness particularly in the auditory and tactile domains may increase sensory cortical input and further contribute to elevated cortical arousal (Halstead et al., 2021; Orefice et al., 2019; Richdale & Schreck, 2009; Smullen et al., 2025). Hyperarousal may be caused by difficulties in autonomic system regulation with increased sympathetic drive at bedtime, yielding elevated heart rate and reduced heart rate variability that may interfere with sleep initiation (Correia et al., 2023; Cosgrave et al., 2021). Cognitive and emotional regulation mechanisms may include intrusive thoughts and rumination that generate anxiety at bedtime and interfere with sleep initiation (Harvey, 2002). Note that these different mechanistic explanations are not mutually exclusive and likely interconnected. Moreover, hyperarousal will interact with basic sleep homeostasis and circadian rhythm mechanisms that govern sleep regulation (Borbély, 2022). For example, hyperarousal may generate behaviors that increase light exposure at night (e.g., screen use), thereby impacting circadian sleep regulation.

The hyperarousal theory of Insomnia may be particularly useful for studying sleep initiation difficulties in autism and ADHD, which co-occur frequently (i.e., 50% of autistic children also have ADHD) and exhibit considerable genetic overlap (Gadow et al., 2006; Rommelse et al., 2010). Studies examining caregiver questionnaires have reported that the individual magnitudes of hyperactivity (Owens, 2008; Thomas et al., 2018), sensory sensitivity (Lane et al., 2022; Manelis-Baram et al., 2021), and anxiety (Galli et al., 2022; Moore et al., 2017) are associated with the severity of sleep disturbances in children with ADHD and/or autism. However, previous studies did not use direct and objective measures of physiological hyperarousal or SOL.

To study sleep initiation difficulties in children with and without developmental conditions we utilized data available in the Simons Sleep Project (SSP) dataset (Hacohen et al., 2025). The SSP contains data recorded with three synchronized devices from >3,600 nights in 102 autistic children and 98 of their siblings. Devices include the Dreem3 EEG headband (Beacon Inc.) that records raw EEG from five channels (Arnal et al., 2020) and the EmbracePlus multi-sensor smartwatch (Empatica Inc.) that records actigraphy, photoplethysmography (PPG), and skin temperature. SSP data is particularly useful for studying sleep because each participant was recorded during multiple nights in their home environment, mitigating potential concerns regarding first night effect (Byun et al., 2019) that are particularly relevant to autism and ADHD research (Primeau et al., 2016). The available data allowed us to quantify changes in neural, physiological, and behavioral measures as participants transitioned from wake to sleep, enabling us to determine whether individual differences in SOL could be explained by difficulties regulating arousal and initiating healthy sleep physiology.

## Methods

### Participants and data collection

Recruitment and socio-demographic details of SSP participants have been published previously (Hacohen et al., 2025). The initial SSP dataset includes sleep recordings of >3,600 nights from 102 autistic children and 98 of their non-autistic siblings, all 10-17-years-old, who previously participated in the Simons Powering Autism Research (SPARK) study (Feliciano et al., 2018). Social Responsiveness Scale, 2^nd^ edition (SRS-2) scores are available for all participants and demonstrate that autistic children had significantly higher autism symptoms than their non-autistic siblings (Table 1). Devices were mailed to participating families throughout the U.S. who recorded sleep in each participant for approximately 3 weeks, on average. However, some of the participants did not record with all the devices due to technical or compliance issues and some recordings were of poor quality and could not be analyzed (see data preprocessing and cleaning sections). In the current study we, therefore, analyzed data from a final sample of 95 autistic children and 91 siblings (Table 1). Of these children, 181 and 169 successfully recorded high quality data with the Dreem3 EEG headband (Beacon Biosignals Inc.) and EmbracePlus smartwatch (Empatica Inc.), respectively, as described below.

**Table 1:** Characteristics of SSP participants who were analyzed in the current study including age, sex, and parent reported SRS scores, medical diagnoses, and medication use as categorized in the SSP (Hacohen et al., 2025). Mean (M), standard deviation (std), number (n), and percentage (%), are reported per measure as relevant.

| Variable | Autism (n=95) | Siblings (n=91) | Statistics |
| --- | --- | --- | --- |
| Age (years) | 13.63 ± 2.10 | 13.24 ± 2.17 | t=1.24, p=0.215 |
| Sex (n, males/females) | 80/15 | 51/40 | $\chi^2=19.73$, p<0.001 |
| SRS-2 total score | 73.60 ± 10.3 | 47.71 ± 7.27 | t=19.86, p < 0.001 |
| **Medical diagnoses** |  |  |  |
| ADHD n(%) | 51 (53.7%) | 29 (31.9%) | $\chi^2=8.16$, p=0.004 |
| Depression | 14 (14.7%) | 10 (11.0%) | $\chi^2=0.30$, $p=0.587$ |
| Anxiety | 30 (31.6%) | 21 (23.1%) | $\chi^2=1.29$, $p=0.256$ |
| **Medication** |  |  |  |
| Any medication | 60 (63.2%) | 45 (49.5%) | $\chi^2=3.02$, $p=0.082$ |
| Antidepressants | 28 (29.5%) | 12 (13.2%) | **$\chi^2=6.37$, $p=0.012$** |
| Stimulants | 28 (29.5%) | 12 (13.2%) | **$\chi^2=6.37$, $p=0.012$** |
| Antianxiety | 22 (23.2%) | 13 (14.3%) | $\chi^2=1.85$, $p=0.174$ |
| Antipsychotics | 8 (8.4%) | 1 (1.1%) | **$\chi^2=4.79$, $p=0.047$** |
| Sleep aids | 8 (8.4%) | 7 (7.7%) | $\chi^2=0.00$, $p=1.000$ |
| Anticonvulsants | 3 (3.2%) | 2 (2.2%) | $\chi^2=0.00$, $p=1.00$ |

### Devices and data types

We analyzed SSP sleep recordings from two devices: the Dreem3 EEG headband and the EmbracePlus smartwatch. The Dreem3 data includes raw EEG recorded from frontal and occipital channels (F1, Fz, F2, O1, O2) at 250 Hz and accelerometer data recorded at 50 Hz, which are available in an EDF file per night. In addition, automated Dreem3 sleep-staging data, a hypnogram, and summary sleep statistics (e.g., TST, WASO) are available in a separate text file per night. The Empatica EmbracePlus smartwatch data includes raw accelerometer and photoplethysmography (PPG) data recorded at 64 Hz and skin temperature recorded at 1 Hz, which are available in 24-h CSV files per sensor. In addition, automated Empatica algorithms derive skin temperature, heart rate, heart-rate variability, movement counts, and sleep–wake classification per minute, which are available in separate, corresponding 24-h CSV files. Further details on data acquisition and structure are provided in the SSP paper (Hacohen et al., 2025).

### Sleep Onset and Sleep Onset Latency (SOL)

Families were instructed to begin the Dreem recording when the participant was ready to go to sleep. We, therefore, computed sleep onset latency (SOL) per night as the time interval between initiation of the Dreem recording and sleep onset (i.e., first sleep epoch) as determined by the Dreem automated sleep-staging algorithm. The Dreem sleep-staging algorithm is based on a long short-term memory (LSTM) neural network that was trained to classify sleep stages in 30 second epochs according to multiple EEG spectral features and accelerometer derived movement features (Arnal et al., 2020). Sleep onset and SOL were successfully identified and computed in 2,534 nights of 194 participants. SOL values were averaged across valid nights, as described below, to compute a mean SOL value per participant.

### Dreem3 EEG data preprocessing and cleaning

Since SSP data was recorded at home by families in uncontrolled settings, we implemented strict data cleaning procedures that ensured our analyses were performed with reliable, high-quality data. Raw EEG recordings from the 2,534 nights described above were preprocessed with custom written code in Python. Each recording was band-pass filtered (0.5-25 Hz) and segmented into consecutive 30-second epochs. We then computed multiple time and frequency domain features per epoch and EEG channel. Time-domain features included variance, kurtosis, mobility, complexity, and maximum absolute amplitude. Note that these features are commonly used to assess signal quality in EEG recordings (Fox et al., 2024). Frequency-domain features included absolute and relative power in the delta (0.75-4 Hz), theta (4.1-8 Hz), alpha (8.1-12Hz), and beta (12.1-16 Hz) frequency bands as well as total power (0.75-25 Hz). Absolute power was calculated per frequency band using the Welch’s method. Relative power was computed by dividing absolute power in each frequency band by total power.

For every subject, we identified and excluded epochs with extreme feature values relative to their distribution in the sleep initiation window from 20 minutes before sleep onset to 40 minutes after, separately for each channel. We defined an exclusion threshold for each measure, per subject, that corresponded to the median plus three times the inter-quartile range as follows:

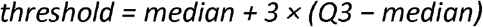

Epochs with kurtosis, variance, maximum amplitude, or root total power that exceeded this threshold were flagged. Nights in which more than 30% of epochs within this window were flagged were excluded from further analysis. Participants with fewer than three valid nights were removed from the dataset. This step resulted in the exclusion of 144 nights from the autism group and 160 nights from the sibling group as well as the removal of 2 autistic children and 4 siblings.

Next, all epochs underwent a second outlier detection step at the sample level. Epochs were flagged if they exceeded the 95th percentile of the entire sample distribution (i.e., across participants) using the same measures: kurtosis, variance, maximum amplitude, and root total power. Nights with more than 30% flagged epochs and participants with fewer than three valid nights were excluded from further analyses. This step excluded an additional 72 nights from the autism group and 88 nights from the sibling group as well as the removal of 4 autistic children and 3 siblings. A total of 480 nights and 13 participants were excluded across both steps, yielding a final EEG dataset with 2054 nights from 181 children.

In all included data, epochs flagged as noisy were replaced with NaN values and linear interpolation was performed to replace the missing values per measure, yielding a time-course of values from 20 minutes before sleep onset to 40 minutes after. We then smoothed each time-course using a 5-minute moving average with a step size of one epoch.

Finally, to remove individual differences in baseline EEG characteristics across participants and night-by-night differences in Dreem3 application, we normalized each EEG measure per night according to its pre-sleep baseline. Specifically, we subtracted the mean value in the 20 minutes preceding sleep onset from the entire time-course. This ensured that we were isolating relative changes associated with sleep onset, applying a similar logic to that of event related potential (ERP) and functional Magnetic Resonance Imaging (fMRI) analyses.

### EmbracePlus data cleaning

EmbracePlus smartwatch recordings were initially available for 2,251 nights from 192 participants who simultaneously recorded their sleep with the Dreem3 headband (i.e., sleep onset and SOL as defined by the Dreem3 were available for these nights). We extracted the Empatica derived movement counts, heart rate (HR), and skin temperature values per minute in the sleep initiation window from 20 minutes before sleep onset to 40 minutes after. The EmbracePlus data also includes an automated wear-detection measure per minute. We flagged 1-minute epochs where this measure was <70% for exclusion. We also flagged epochs with HR values lower than 40 and higher than 200 bpm as well as skin temperature below 20°C or above 40°C. Nights in which more than 30% of epochs in the sleep initiation window were flagged were excluded from further analysis. Participants with fewer than three valid nights were also excluded from further analysis. This step excluded 202 nights from the autism group and 152 nights from the sibling group along with 12 autistic children and 11 siblings, yielding a final EmbracePlus dataset of 1,897 nights from 169 participants.

In all included data, epochs flagged as noisy were replaced with NaN values and linear interpolation was performed to replace the missing values per measure, yielding a time-course of values from 20 minutes before sleep onset to 40 minutes after. We then smoothed each time-course using a 5-minute moving average with a step size of one epoch.

Here too, we removed individual differences in baseline HR, skin temperature, and movement counts as well as night-by-night differences in EmbracePlus application by normalizing each measure per night according to its pre-sleep baseline. Specifically, we subtracted the mean value in the 20 minutes preceding sleep onset from the entire time-course.

### Analysis of transitions at sleep onset

To capture the timing, speed, and magnitude of neural, physiological, and behavioral transitions associated with sleep onset, we fit each measure (i.e., delta power, skin temperature, heart rate, and movement count) with the following sigmoid function:

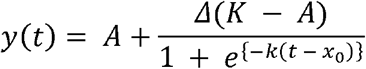

Where, y(t) is a time-course vector of the measure from 20 minutes before sleep onset to 40 minutes after sleep onset, A is the left asymptote of the sigmoid, K is right asymptote of the sigmoid (K-A is the magnitude of transition), x0 is the inflection point of the sigmoid (timing of the transition), and k is the slope of the sigmoid (velocity of the transition). Using custom written Python code and the curve_fit function from the scipy.optimize library, we fit the sigmoid function to the mean data across nights per participant (Figure 1A). The fitting procedure required setting initial values and bounds for the parameters. The lower and upper asymptote values were set as the 5^th^ percentile of the pre-sleep segment and 95^th^ percentile of the post-sleep segment, respectively. Bounds were set to ±50% around these values to stabilize the fits. The inflection point (x0) was initialized as the time with the steepest slope within the sleep initiation window, while the initial slope parameter (k) was defined as 4 divided by the range of the sleep initiation window and bounded between −50 and +50 (Figure 1A).

**Figure 1:**
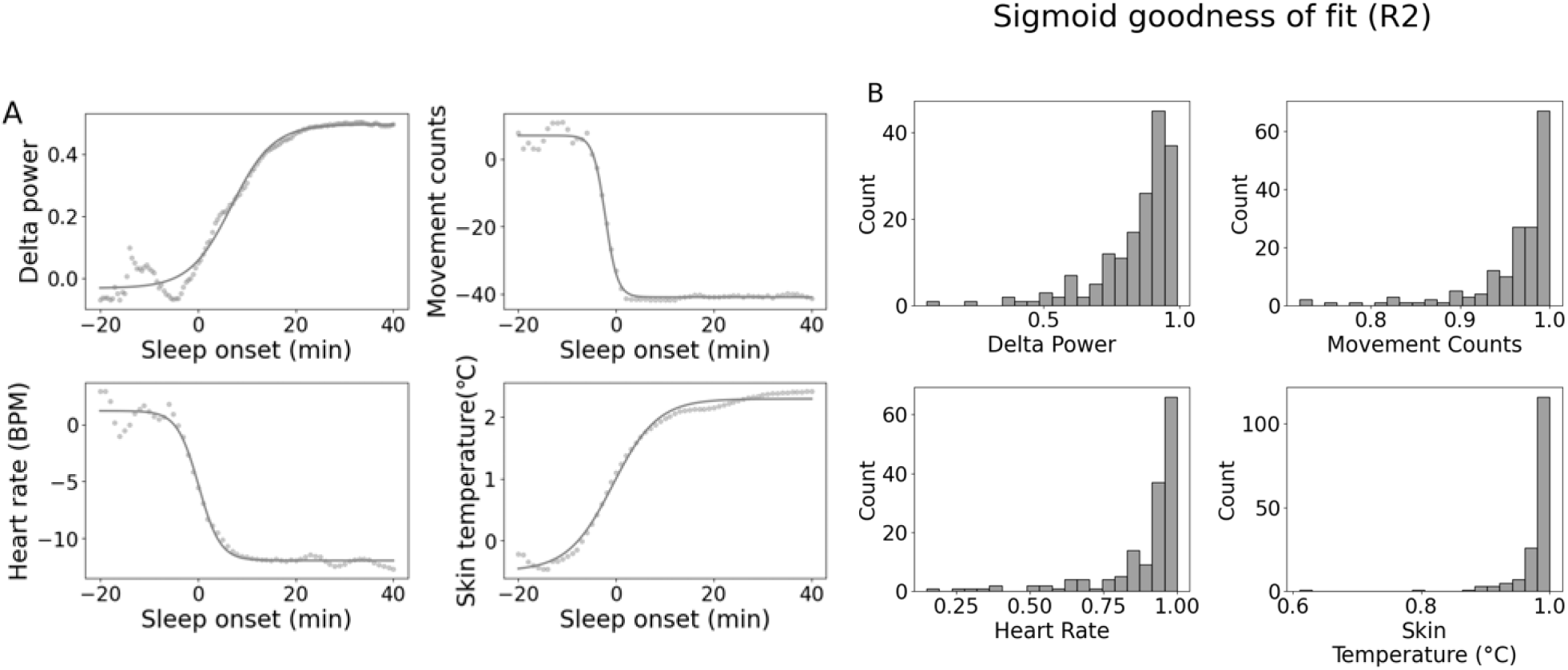
Sigmoid fit per measure. **A**: Example of sigmoid fits for one participant using the mean data (across nights) from each of four measures examined in the study. Smoothed data (dotted line) are overlayed with fitted sigmoid function (solid line) in the sleep initiation window from 20 minutes before sleep onset to 40 minutes after. **B**: Histograms of goodness-of-fit (R^2^) values from all participants per measure.

Sigmoid fits were successfully computed for relative delta power in 173 participants (out of 181 participants with valid EEG data) and for skin temperature, movement counts, and heart rate in 163, 168, and 155 participants, respectively (out of 169 participants with valid EmbracePlus data). Sigmoid fits for all four measures were available in 141 participants (71 autism, 70 siblings). The sigmoid fits were excellent for all measures (Figure 1B), with median R^2^ values of 0.98 for movement counts (IQR = 0.04), 0.99 for skin temperature (IQR = 0.02), 0.89 for EEG delta power (IQR = 0.16), and 0.95 for heart rate (IQR = 0.1). An equivalent night-wise analysis, in which sigmoid models were fit separately to data from individual nights, yielded similar results across all measures (Supplementary Figure 1S; Table 1S).

This analysis yielded three important measures characterizing the transition from wake to sleep for each measure. The first was transition magnitude, which was estimated as the difference across Sigmoid asymptotes (K - A). The second was transition timing, which was estimated as the inflection point (x0) of the Sigmoid. The third was transition velocity, which was estimated as the slope (k) of the Sigmoid.

### Statistical analysis

All analyses were performed with custom written code in Python using the scipy, statsmodels, and pandas toolboxes. We performed linear mixed-effects linear model (MLM) analyses to assess the relationship between the children’s SOL lengths and transitions in each of the four measures (i.e., delta power, hear rate, movement counts, and skin temperature). We created a separate MLM for transition magnitudes (K - A), transition timing (x0), and transition speed (k), using the relevant Sigmoid function parameters. We included the four measures in each model and examined which transition parameter (magnitude, timing, or speed) was most strongly associated with SOL per participant, thereby attempting to explain SOL differences across participants. Individual subjects were, therefore, defined as fixed effects and family identity was defined as the random effect. Age, sex, and goodness-of-fit (R^2^) of the Sigmoid per subject were also included as fixed effects (i.e., covariates) to ensure that these sources of variability did not impact our results. We compared the marginal R^2^ of these models to assess their relative performance (Nakagawa & Schielzeth, 2013).

To assess whether each MLM explained a significant proportion of SOL variance, above chance level, we conducted a permutation/randomization analysis with 5000 iterations. In each iteration we randomly permuted SOL values across participants, thereby breaking the true association between predictors and outcomes while preserving the overall distribution of the data. We computed the marginal R^2^ value for each iteration, generating a null distribution of results expected by chance. We then calculated the percentile of the true marginal R^2^ value relative to the null distribution. The p-value was calculated as (*k* + 1)/(*B* + 1), where k is the number of permuted R^2^ values greater than or equal to the observed R^2^ and *B* is the total number of permutations.

In additional analyses, we split participants into four groups according to their SOL values with each group corresponding to a quartile of the SOL distribution. We then assessed whether transition magnitudes in delta power, movement counts, skin temperature, or heart rate differed significantly across the four SOL groups using MLM analyses with age, sex, and Sigmoid goodness of fit (r^2^) included as covariates. Additional post-hoc analyses using specific linear model contrasts were used to determine whether there were significant differences across specific SOL subgroups.

Finally, we split the children into four groups according to their diagnostic classification: autism, ADHD, autism+ADHD, and typical development. We performed analogous MLM analyses to compare transition magnitudes in delta power, movement counts, heart rate, and temperature across the four diagnostic groups while including age, sex and Sigmoid goodness of fit (r^2^) as covariates. We also compared SOL across the same groups. Individual subjects were modeled as fixed effects and family identity was defined as the random effect. Specific linear model contrasts were used to compare typically developing children with children who had any developmental condition (autism, ADHD, or autism+ADHD). Statistical significance was set to α<0.05 in all analyses.

## Results

Analyses were performed with a final sample of 186 children, which included 95 autistic children and 91 of their siblings (Table 1). We focused our analysis on the sleep initiation period that was defined from 20 minutes before sleep onset to 40 minutes after sleep onset. Sleep onset was identified by the Dreem3 automated sleep-staging algorithm. We extracted corresponding time-courses of EEG delta power, movement counts, heart rate, and skin temperature within this pre-defined period (see Methods).

We split participants into four groups corresponding to four quartiles of the SOL distribution. We then compared sleep initiation time-courses across the four quartiles (Figure 2). This analysis revealed dramatic transitions in delta power, movement counts, heart rate, and skin temperature within the sleep initiation window. Most importantly, the timing and speed (slope) of these transitions was similar across SOL groups, yet the magnitude of transitions was larger in participants who fell asleep quickly (first quartile) versus those who fell asleep slowly (fourth quartile).

**Figure 2:**
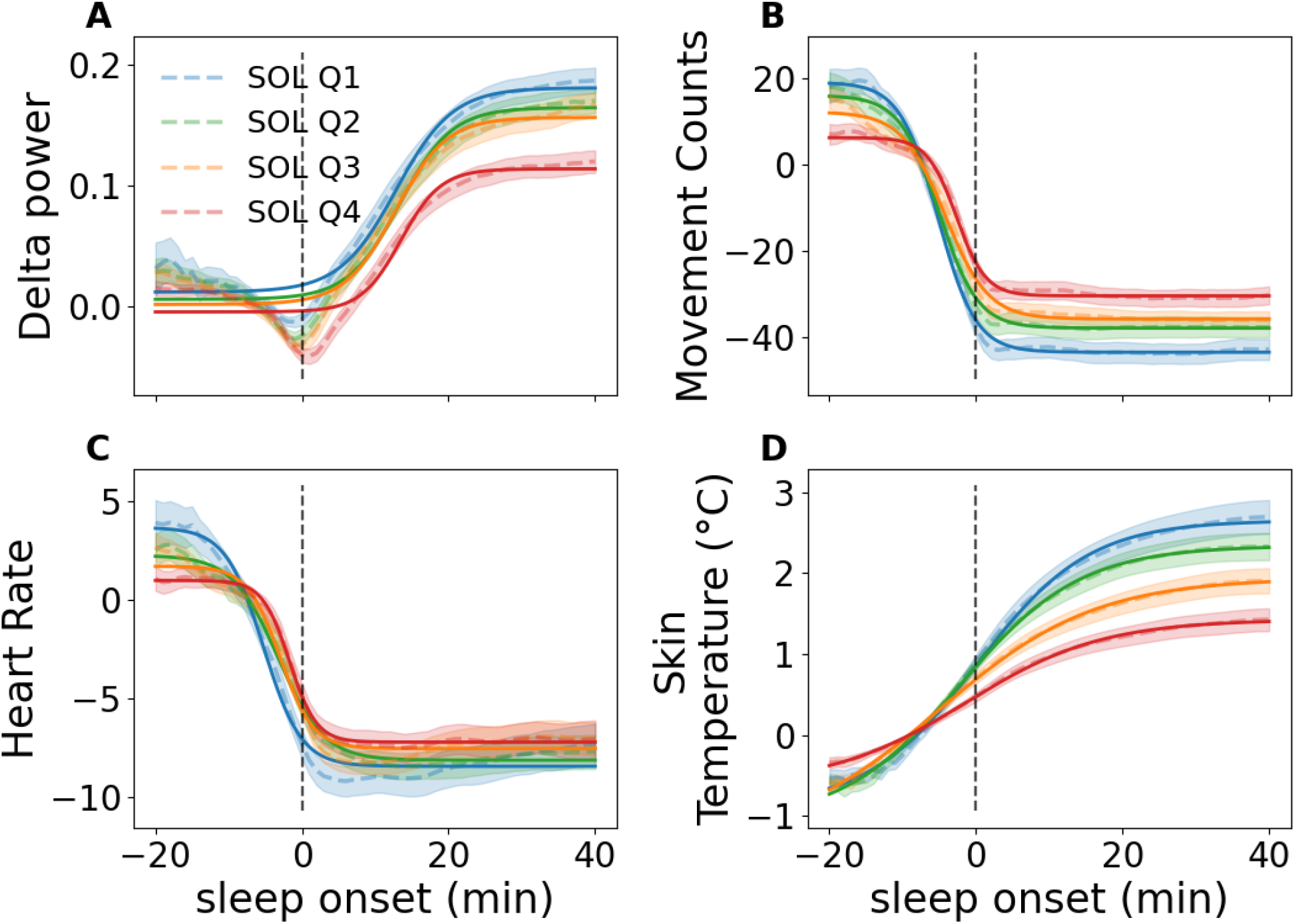
Comparison of sleep onset transitions in four groups of participants corresponding to their SOL quartiles: blue (1^st^ quartile, shortest SOL), green (2^nd^ quartile), orange (3^rd^ quartile), and red (4^th^ quartile, longest SOL). All values are normalized to the pre-sleep period (−20 min to sleep onset). **A**. Relative Delta power **B**. Movement counts **C**. Heart rate **D**. Skin temperature. **Dashed lines**: mean time-courses across participants. **Shaded areas**: ±2 standard error of the mean (SEM). **Solid lines**: Sigmoid fit. **Horizontal dashed line**: Sleep onset according to Dreem3.

We fit a Sigmoid function to the mean data (across nights) of each participant (Figure 1) to estimate the timing, speed, and magnitude of sleep onset transition per measure. This was followed by MLM analyses where we assessed whether the Sigmoid parameter estimates (e.g., transition magnitude) could explain differences in SOL across participants. We created a separate MLM for each of the three Sigmoid parameters: inflection point (x0) indicative of transition timing, slope (k) indicative of transition speed, and difference across asymptotes (K– A) indicative of transition magnitude. Each MLM included four predictors corresponding to the four examined measures (EEG delta power, movement counts, HR, and skin temperature) as well as sex, age, and goodness of fit (r^2^).

Remarkably, the MLM with estimated transition magnitudes from the four measures explained over 25% of the SOL variability across participants, which was more than twice the variability explained by MLMs with transition speed (k) or timing (x0). This demonstrated that transition magnitudes are key to explaining SOL differences across participants.

We performed a permutation analysis to determine whether each model explained significantly more variance than expected by chance (see methods) and found that only the transition magnitude MLM yielded significant result (R^2^_m_ = 0.262; p = 0.009). Within this model EEG delta power (β = −28.25, SE = 9.23, p = .002), skin temperature (β = −2.115, SE = 0.83, p = .011), and movement count (β = −0.31, SE = 0.08, p < .001) predictors were significantly associated with SOL, but HR was not (β = −0.011, SE = 0.26, p = .96). Note that transition magnitudes in movement counts were strongly correlated with HR (r= 0.45, p < 0.001), but not with EEG delta power (r= 0.03, p = 0.69) or temperature (r= −.12, p= 0.15). Hence, HR and movement count transitions may reflect a common phenomenon and may be used interchangeably to explain SOL (see Discussion).

Sex, age, and sigmoid goodness-of-fit (r^2^) predictors within the model described above were not significantly associated with SOL. The MLMs with Sigmoid parameters for transition speed (R^2^_m_= 0.105; p = 0.3 or timing (marginal; R^2^_m_ = 0.126; p = 0.203) did not explain a significant proportion of the variability.

In complementary MLM analyses we assessed differences across children in the four SOL quartile groups (Figure 4). We built a separate MLM for each measure (delta power, movement counts, skin temperature, and heart rate) to determine the ability of transition magnitudes in each measure to explain SOL differences across SOL sub-groups. Age, sex, and goodness-of-fit (r^2^) were included as well to ensure that potential differences across sub-groups were not driven by these factors. Significant differences across SOL sub-groups were apparent in MLMs of all measures. Transition magnitudes differed between the shortest SOL group (Q1) and the longest SOL group (Q4) for skin temperature (R^2^_m_ = 0.179; Q4 vs Q1: β = −1.47, p < .001), heart rate (R^2^_m_ = 0.195; Q4 vs Q1: β = −4.57, p = .009), movement counts (R^2^_m_ = 0.274; Q4 vs Q1: β =−22.16, p < .001), and delta power (R^2^_m_ = 0.289; Q4 vs Q1: β = −0.125, p < .001). In all cases, longer SOL was associated with smaller/weaker transition magnitudes relative to shorter SOL.

Next, we compared SOL and transition magnitudes per physiological/behavioral measure across children with autism (n=43), ADHD (n=28), autism+ADHD (n=50), and typically developing controls (n=50). These sub-groups were defined according to parent reported clinical diagnoses and usage of medication as available in the SSP. Children were included in the control group if they had no neurodevelopmental, psychiatric, or neurological disorders and were not treated with any psychiatric, stimulant, or anti-seizure medicine. Children with alternative diagnoses (e.g., siblings with anxiety and/or depression) were excluded. Estimates of transition magnitude for delta power, movement counts, heart rate, and skin temperature were available in 154, 151, 140, and 146 of the children in this sub-sample, respectively.

We built a MLM for each physiological/behavioral measure and examined whether transition magnitudes differed across diagnostic groups while including age, sex, and goodness-of-fit as covariates (Figure 5). The results revealed that transition magnitudes were generally smaller in the clinical groups than the control group. Transition magnitudes in EEG delta power were smaller in children with autism, ADHD, or both, relative to controls (autism: β = −0.65, p = 0.029;ADHD: β = −0.074, p = 0.027; autism+ADHD: β = −0.06, p = 0.048; R^2^_m_ = 0.24). A linear model contrast comparing control children to the three clinical groups combined revealed a significant difference (β = 0.066, z = −2.68, p = 0.007).

Similarly, transition magnitudes in movement counts were significantly smaller in autism and autism+ADHD relative to control children (autism: β = −15.01, p = 0.001; R^2^_m_ = 0.201; autism+ADHD: β = −12.59, p = 0.011), but the ADHD group did not show the same effect (β =−6.78, p = 0.221). The linear model contrast comparing control children to the three clinical groups revealed a significant difference (β = 11.46, z = 2.89, p = 0.004). Transition magnitudes in heart rate were smaller in the autism group relative to control children (β = −3.18, p = 0.047; R^2^_m_ = 0.169), with a similar non-significant trend for autism+ADHD children (β = −3.25, p = 0.072), but not for ADHD (β = −1.67, p = 0.376). The linear model contrast comparing control children to the three clinical groups revealed a marginally significant difference (β = 2.7, z =1.95, p = 0.052). Finally, transition magnitudes in skin temperature did not differ across diagnostic groups (all group p-values > 0.4; R^2^_m_ = 0.113).

A corresponding MLM analysis demonstrated that SOL was significantly longer in children with autism, ADHD, or both (autism: β = 10.75, p = 0.018; ADHD: β = 14.20, p = 0.005; autism +ADHD: β = 14.51, p = 0.002). A linear model contrast comparing control children to the three clinical groups revealed a highly significant difference (β = −11.9, z = −3.67, p < 0.001).

## Discussion

Falling asleep is a complex process that requires appropriate regulation of multiple interacting neural and physiological systems that transition from a state of wakefulness to sleep (Kräuchi & Wirz-Justice, 2001b; Ogilvie, 2001; Wyatt et al., 1999; Zulley et al., 1981). Our results reveal that individuals with difficulties falling asleep (i.e., long SOL) exhibit weak neural, physiological, and behavioral transitions from wake to sleep that are evident in EEG delta power, HR, skin temperature, and movement count measures (Figure 2). Remarkably, a model with transition-magnitude estimates of these four measures explained more than a quarter of the variance in SOL values across participants, while models with transition speed or timing estimates explained less than half of this variance (Figure 3). A central finding of this study, therefore, is that the magnitude of described neural, physiological, and behavioral transitions is key to explaining sleep initiation problems.

**Figure 3:**
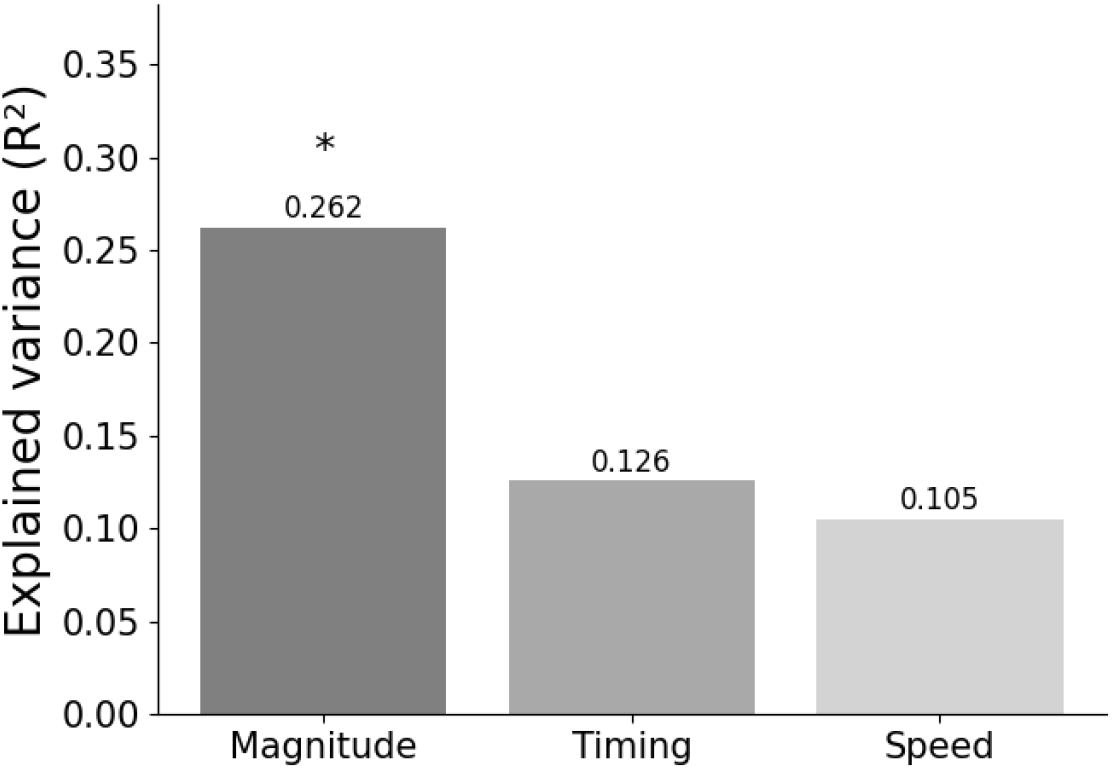
Comparison of three MLMs explaining SOL differences across participants according to estimated EEG delta power, movement counts, heart rate, and skin temperature transitions. **Left**: MLM with transition magnitude estimates. **Middle**: MLM with transition timing estimates. **Right**: MLM with transition speed/slope estimates. Transition magnitude, speed, and timing was estimated per participant by fitting a Sigmoid function to their mean data (across nights). Significance of the models was assessed with a permutation/randomization test (see Methods). **Asterisk**: MLM explained a significant proportion of variance in SOL across participants (p<0.01).

A complementary analysis comparing EEG delta power, HR, skin temperature, and movement count transitions across children in four SOL quartile groups also demonstrated consistent differences with fast sleepers exhibiting significantly larger transitions than slow sleepers (Figure 4). Hence, difficulties falling asleep are associated with poor regulation of multiple neural and physiological mechanisms that are central to the sleep initiation process. These mechanisms seem to be particularly dysregulated in children with autism and/or ADHD (Figure 5).

**Figure 4:**
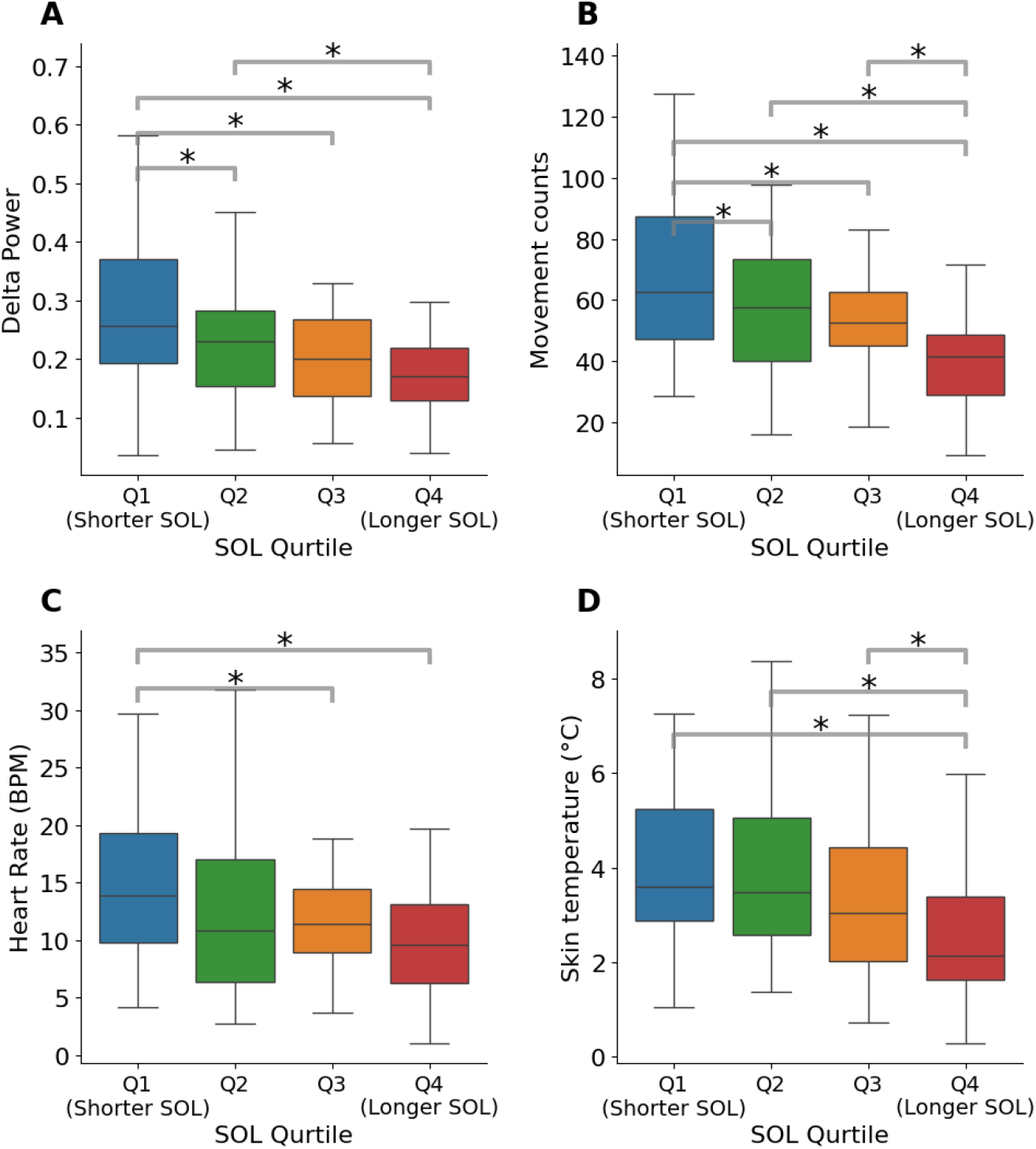
Boxplots show transition magnitudes in EEG delta power (**A**), movement counts (**B**), heart rate (**C**), and skin temperature (**D**) across SOL groups. Linear MLM analyses controlling for age and sex revealed significant differences between participants in the four SOL quartile groups such that participants with longer SOL exhibited weaker transitions in each of the four examined measures. **Asterisks**: Significant differences across specific SOL groups (p<0.05) as determined with a contrast test.

**Figure 5.**
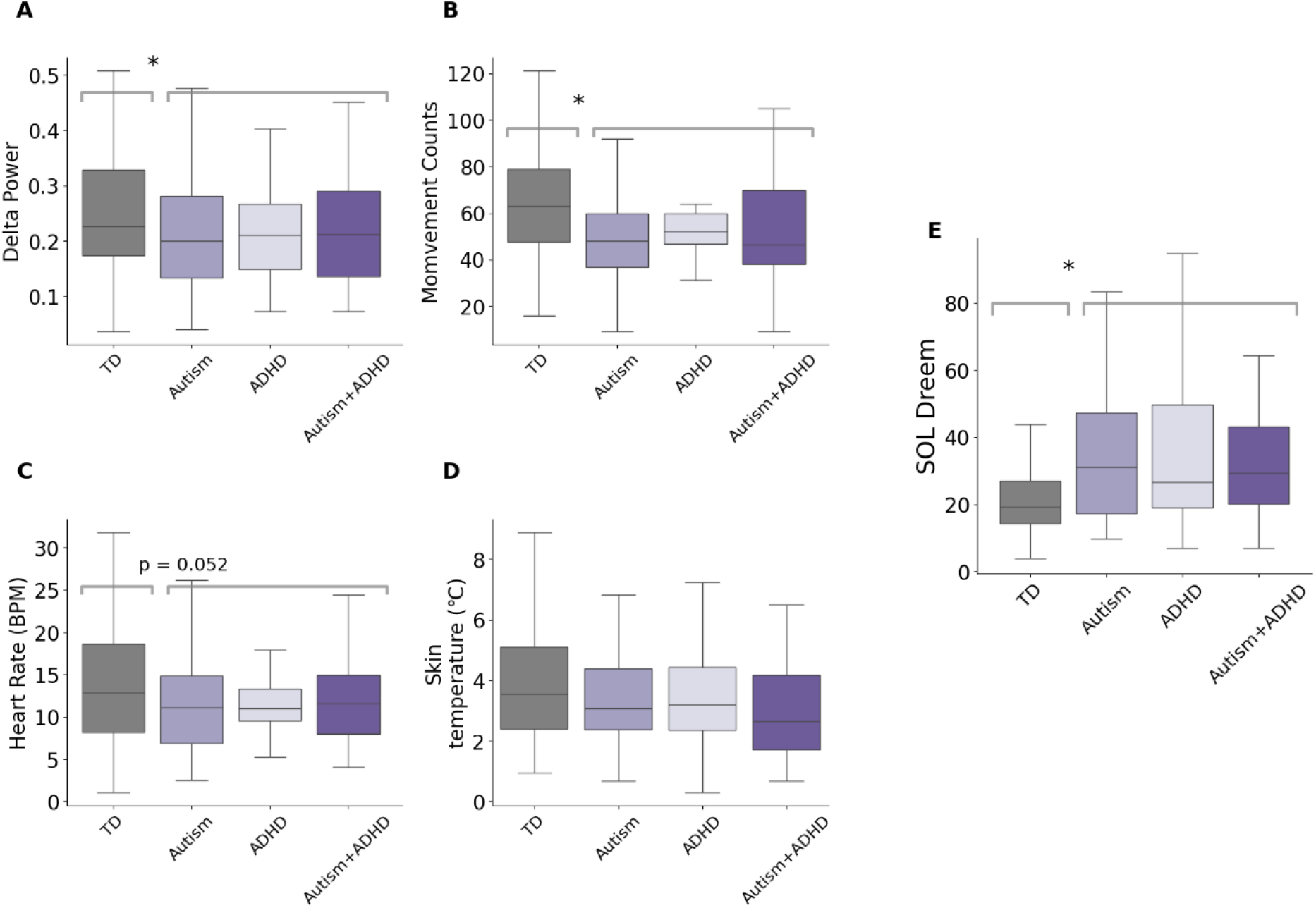
Comparison of transition magnitudes in EEG Delta Power (**A**), Movement Counts (**B**), Heart Rate (**C**), and skin temperature (**D**) as well as a comparison of SOL (**E**) across children with typical development (gray), autism (purple), ADHD (light purple), and both autism and ADHD (dark purple). **Asterisk**: significant difference in a linear contrast analysis between control children and the three clinical groups (p<0.05).

### Multiple dysregulated mechanisms may prolong SOL

Previous research has thoroughly characterized neural transitions in the central nervous system of humans as they fall asleep (Ogilvie, 2001). While the epoch of sleep onset is formally defined by visual identification of subtle changes in EEG power according to AASM rules (Troester et al., 2023), initiation of stable sleep is a lengthier process characterized by a gradual increase in delta power (i.e., SWA) that grows with the depth of sleep (Borb & Achermann, 1999; Dijk & Von Schantz, 2005; Esser et al., 2007; Magnin et al., 2010). Many studies have demonstrated that the magnitude of SWA increases with the extent of sleep deprivation, making SWA a potent measure of homeostatic sleep drive or sleep pressure (Achermann et al., 1993; Borb & Achermann, 1999; Borbély, 2022; Dijk & Von Schantz, 2005). To study the transition into stable sleep in the current study we examined changes in delta power from 20 minutes before to 40 minutes after sleep onset. Our results reveal that children with difficulties falling asleep exhibit weaker SWA transitions with smaller SWA amplitude increases relative to their pre-sleep period. This could reflect poor homeostatic sleep drive, as reported previously in autism (Arazi et al., 2020), or the existence of cortical hyperarousal (Riedner et al., 2016; Spiegelhalder et al., 2012), that may interfere with the initiation of SWA, slowing down the transition into stable deep sleep and potentially extending SOL.

Changes in thermal regulation also play an important part in sleep initiation. Numerous studies have demonstrated that the circadian evening release of Melatonin triggers peripheral vasodilation in the hands and feet (Kräuchi et al., 2006), which causes an increase in skin temperature that facilitates heat loss, reducing core body temperatures, and increasing the propensity to sleep (Kräuchi & Wirz-Justice, 2001; Shochat et al., 1998; Zulley et al., 1981). Poor skin temperature transitions in children with longer SOL suggest potential dysregulation of Melatonin release, a mismatch in the timing of sleep onset relative to the endogenous circadian rhythm, or dysregulation of vasodilation or thermal regulation. Alternatively, hyperarousal may drive behaviors that alter light exposure (e.g., use of screens), thereby interfering with proper circadian control of thermoregulation. While there may be multiple underlying causes for thermal dysregulation at sleep onset, our results suggest that this dysregulation is associated with prolonged SOL.

Heart rate is another key physiological measure that decreases noticeably as humans transition from wake to sleep (De Zambotti et al., 2011; Kräuchi & Wirz-Justice, 2001b; Shinar et al., 2006). This transition is regulated by autonomic nervous system changes with increased parasympathetic drive, which lowers HR and shifts multiple physiological systems to a state of relaxation (Tobaldini et al., 2013). The reduction in HR and increase in heart rate variability (HRV) reflect a coordinated downregulation of arousal that facilitates the transition to sleep (Chouchou & Desseilles, 2014; Ma et al., 2024). In individuals with insomnia, attenuated reductions in HR and persistent sympathetic activation have been interpreted as markers of autonomic/physiological hyperarousal (Cosgrave et al., 2021; De Zambotti et al., 2011). In the current study, children with longer SOL exhibited weaker HR transitions (Figure 2C), suggesting that they have difficulty switching from sympathetic to parasympathetic activation, which is necessary for initiating stable sleep. Lower HR before sleep onset in children with long SOL suggests that they were trying to fall asleep but were unable to transition into sleep. Faster HR after sleep onset suggests that children with long SOL remain more physiologically aroused after they finally do fall asleep.

Cessation of movement is a fundamental behavioral marker of the transition from wake to sleep. Indeed, in many animal models, including fish and flies, movement is the main criterion used to define sleep and wake periods (PJ et al., 2000; Yokogawa et al., 2007). In humans, accelerometry has been used for over four decades to estimate sleep–wake patterns in both research and clinical settings (Patterson et al., 2023), identifying sleep periods by low movement counts (Ogilvie, 2001; Sadeh et al., 2000). In the current study, children with longer SOL exhibited smaller reductions in movement counts during the sleep initiation window. Children with longer SOL moved less before sleep onset and more after sleep onset (Figure 2), suggesting that they have difficulty switching from active states to complete relaxation when initiating sleep. Lower movement counts before sleep onset in children with longer SOL suggests that they are trying to fall asleep (i.e., are moving less) but are unable to initiate sleep. Higher movement counts after sleep onset suggest that children with longer SOL remain restless even after they finally do fall asleep. These findings are very similar to those described above for HR. Indeed, movement count and HR transitions were significantly correlated (r= 0.45; Supplementary material Figure 2S), demonstrating that they are inter-related, potentially interfering with sleep initiation and prolonging SOL in tandem.

### Hyperarousal and prolonged SOL

Hyperarousal may provide a unifying framework for interpreting the neural, physiological, and behavioral findings described above. The hyperarousal theory of insomnia posits that difficulties initiating sleep stem from a combination of cortical, physiological, and cognitive arousal components (Riemann et al., 2010). Previous studies have interpreted higher beta/gamma EEG power and lower delta power during non-REM sleep as evidence for cortical hyperarousal, which may represent a difficulty disengaging cortical sensory systems from thalamic input (Levenson et al., 2015; Riemann et al., 2010). Similarly, previous studies have interpreted elevated HR and reduced HRV during sleep as evidence of physiological hyperarousal and increased sympathetic drive that is likely to interfere with sleep initiation (Correia et al., 2023; Cosgrave et al., 2021). Cognitive-emotional hyperarousal including intrusive thoughts, anxiety, and rumination, which are common in children with autism and/or ADHD (see below) may further increase arousal and delay sleep initiation (Carney et al., 2010; Kalmbach et al., 2020). These components likely interact with each other and with circadian and homeostatic sleep regulation mechanisms (Borbély, 2022). For example, cognitive-emotional hyperarousal may increase screen use and light exposure, thereby impacting cortical arousal and circadian sleep regulation. While we did not measure cognitive-emotional hyperarousal in the current study, our results do suggest that children with longer SOL exhibit evidence of cortical and physiological hyperarousal during their sleep initiation period.

### SOL in children with neurodevelopmental conditions

Previous research with the SSP has already demonstrated that autistic children exhibit significantly longer SOL relative to their non-autistic siblings (Hacohen et al., 2025). Here we demonstrate that not only children with autism exhibit prolonged SOL, but also children with ADHD or both conditions exhibit significantly longer SOL relative to control children without any neurological or psychiatric disorders (Figure 5). This is in line with previous research demonstrating that children with ADHD also exhibit severe sleep disturbances (Gruber et al., 2009; Xian et al., 2025). Note that ADHD and autism are closely related disorders where approximately 50% of autistic children also exhibit ADHD symptoms (Ronald et al., 2008).

Most importantly, children with autism and/or ADHD exhibited weaker sleep transitions in most measures when compared to typically developing controls (Figure 5). This suggests that children with autism and/or ADHD exhibit cortical, physiological, and behavioral hyperarousal as previously suggested by others (Baker et al., 2019; Furrer et al., 2019). Difficulties disengaging from stimulating activities, sensory hypersensitivities, bed-time anxiety, and emotional dysregulation are particularly common in autism (Halstead et al., 2021; Manelis-Baram et al., 2021; Mazurek & Petroski, 2015) and ADHD (Tong et al., 2018; Wajszilber et al., 2018), and are likely to exacerbate hyperarousal, weaken sleep transitions, and prolong SOL.

Reduced sleep pressure, previously reported in autism (Arazi et al., 2020), may also be associated with cortical hyperarousal.

### Studying sleep onset with ecological data

A key advantage of the current study was the utilization of multi-night recordings from multiple devices/sensors able to capture neural, physiological, and behavioral data during the sleep initiation period within the children’s natural home environment. This is important for estimating habitual sleep physiology and behavior in an ecological manner and accounting for night-to-night variability. While laboratory PSG recordings are considered the gold standard for sleep research, they typically include only a single night of recording in an unfamiliar and often uncomfortable environment that is known to generate the first⍰night effect (Newell et al., 2012). Indeed, the sleep recorded during such a night is unlikely to represent the child’s habitual sleep at home and this concern is particularly potent when studying children with neurodevelopmental conditions, who are highly sensitive to environmental changes (Bessey et al., 2013; Ezedinma et al., 2025; Lanzlinger et al., 2023). Hence, the SSP offers a unique opportunity to study the sleep initiation period in >2000 nights from children with and without neurodevelopmental conditions using a highly valuable ecological design.

### Limitations

Several limitations should be considered when interpreting the presented findings. First, while home recordings are highly ecological, they are performed in an uncontrolled manner that introduces variability in sensor placement, environmental noise, and data quality. For this reason, we applied strict quality control criteria when including recordings in our analysis. Nevertheless, a potential limitation of SSP data is that it is likely to be of poorer quality than laboratory PSG recordings. Second, our analyses focused on the wake-to-sleep transition window (20 minutes before to 40 minutes after sleep onset) and did not examine physiological and behavioral data from earlier parts of the evening that may have contributed to difficulties falling asleep, such as vigorous physical activity. Finally, differences across diagnostic groups should be interpreted with caution due to the relatively small sample size and the fact that diagnostic labeling in the SSP is based on parent reported clinical diagnosis, which may not be entirely accurate.

## Conclusions

Using extensive ecological data acquired at home with synchronized recordings from multiple devices, our results demonstrate that children with longer SOL exhibit weaker neural, physiological, and behavioral sleep transitions. These findings suggest that multiple dysregulated mechanisms generate hyperarousal in children with sleep initiation difficulties, particularly in children with autism and/or ADHD. Importantly it is the magnitude of transitions that seems to explain differences in SOL across participants rather than the speed or timing of transitions within the sleep initiation period. These findings focus future basic and clinical research into the underlying mechanisms of sleep initiation difficulties and demonstrate the unique value of the SSP dataset for studying sleep disturbances.

## Acknowledgements

This work was supported by the Simons Foundation Autism Research Initiative (SFARI; Award No. 00010173).

