## Supplemental material for "Sleep initiation difficulties involve weaker neural and physiological sleep transitions, particularly in children with neurodevelopmental conditions"

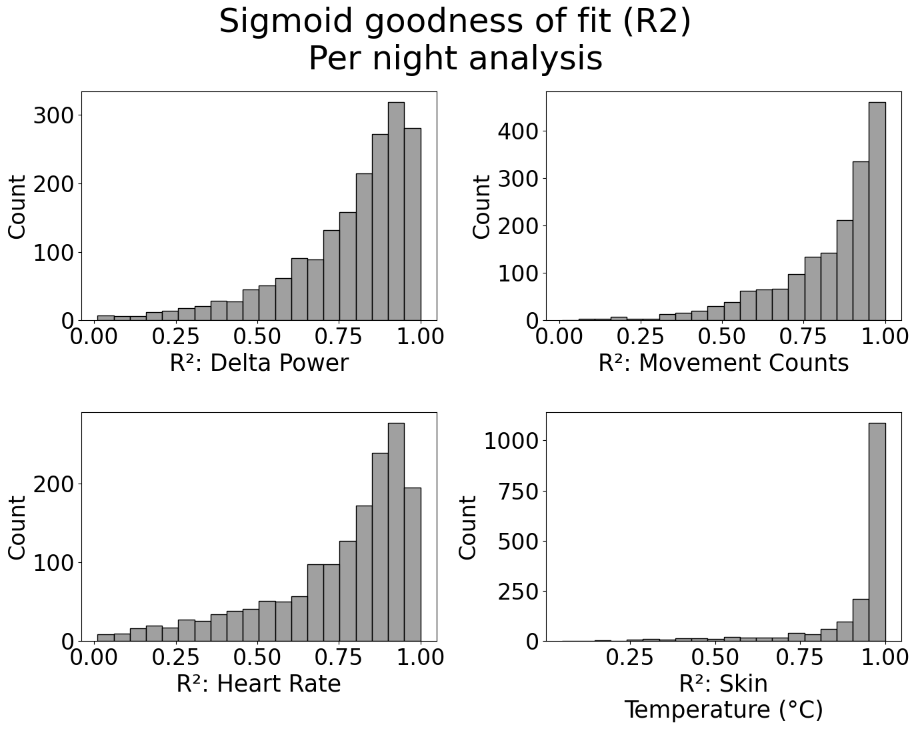

**Figure 1S**: Histograms of goodness-of-fit (R²) values per individual night as computed for each of the following measures. **A.** EEG Delta power. **B.** Movement counts. **C.** Heart rate. **D.** Skin temperature.

| **Measure** | **Median R²** | **IQR R²** | **Q1** | **Q3** |
| --- | --- | --- | --- | --- |
| **EEG Delta Power** | 0.84 | 0.23 | 0.70 | 0.93 |
| **Skin Temperature** | 0.98 | 0.07 | 0.92 | 0.99 |
| **Heart Rate** | 0.83 | 0.26 | 0.66 | 0.92 |
| **Movement counts** | 0.89 | 0.21 | 0.75 | 0.96 |

**Table 1S:** Summary statistics of goodness-of-fit (R²) values per night as estimated for each of the four examined measures.

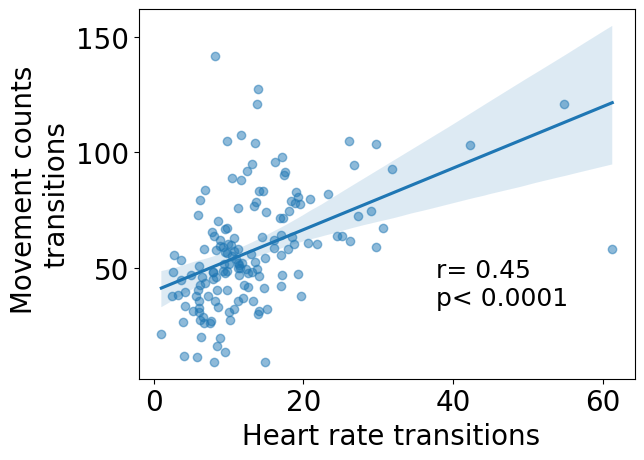

**Figure 2S**: Relationship between movement counts and heart rate transition magnitudes. Each point represents an individual child. Pearson correlation and statistical significance are noted
